# Mutations in an intracellular vestibule that bifurcates from the pore of ClC-1 chloride ion channels affect anion permeation

**DOI:** 10.1101/2020.11.01.364042

**Authors:** Brett Bennetts, Craig J. Morton, Michael W. Parker

**Author notes:** Corresponding Author: Brett Bennetts. Department of Biochemistry and Molecular Biology, Bio21 Molecular Science and Biotechnology Institute, The University of Melbourne, Parkville, Victoria 3010, Australia.

## Abstract

The ubiquitous CLC protein superfamily consists of channels, that permit passive diffusion of Cl ions across biological membranes, and pumps, that can actively transport Cl ions against their electrochemical gradient; yet, puzzlingly, both types share a strongly conserved Cl ion transport pathway comprised of three consecutive binding sites. This raises the question; how does the same pathway support passive diffusion in CLC channels and active transport in CLC pumps? Based on high-resolution structural data current theories suggest that subtle structural differences in the conserved pathway allow CLC channels to ‘leak’ Cl ions. A recent cryo-electron microscopy structure of the human ClC-1 channel does not show occupancy of the central Cl ion binding site but reveals a wide intracellular vestibule that bifurcates from the conserved pathway in this region. Here we show that replacing residues that line the ClC-1 intracellular vestibule with the corresponding residues of CLC pumps resulted in interactions between permeating anions at neighbouring binding sites and altered anion selectivity. Removing the side chain of a strictly conserved tyrosine residue, that coordinates Cl ion at the central binding site of CLC pumps, removed multi-ion behaviour in ClC-1 mutants. In contrast, removing the side chain of a highly conserved glutamate residue that transiently occupies Cl ion binding sites, as part of the transport mechanism of CLC pumps and the mechanism that opens and closes CLC channels, only partially removed multi-ion behaviour in ClC-1 mutants. Our findings show that structural differences between CLC channels and pumps, outside of the conserved Cl ion transport pathway, fundamentally affect anion permeation in ClC-1 channels.

**Summary:** Some CLC proteins are passive Cl^-^ channels while others are active Cl^-^ pumps but, paradoxically, both share a conserved, canonical, Cl^-^ permeation pathway. Here Bennetts, Morton and Parker show that ‘pump-like’ mutations in a poorly conserved region, located remotely from the canonical pathway, affect anion permeation in human ClC-1 channels.

## Introduction

In humans nine members of the ubiquitous CLC superfamily^*a*^ of Cl ion transporting proteins perform vital physiological functions that are underscored by mutations associated with a wide range of human diseases (Jentsch and Pusch, 2018). Despite sharing remarkably similar structures, the CLC superfamily is divided into dissipative Cl^-^ channels (four in humans) and active Cl^-^ pumps (five in humans). It is currently unclear how such closely related proteins accomplish functions that, on thermodynamic considerations alone, are seemingly incompatible (Miller, 2006). CLC ion channels permit diffusion of Cl^-^ down its electrochemical gradient while CLC pumps, commonly known as exchangers, harness energy evolved from dissipation of an H^+^ gradient to power pumping of Cl^-^ against its gradient, or *vice versa* (Jentsch and Pusch, 2018). Surprisingly, despite the disparate functions of CLC channels and exchangers, recent cryo-EM structures show that both functional classes share a strongly conserved Cl^-^ transport pathway (Park et al., 2017; Park and MacKinnon, 2018).

The Cl^-^ transport pathway of CLC exchangers is comprised of three consecutive binding sites (Dutzler et al., 2002; Dutzler et al., 2003; Feng et al., 2010), typically referred to as *S*_ext_, *S*_cen_, and *S*_int_ according to their proximity to the extracellular (ext) or intracellular (int) aqueous solutions. All three binding sites may be simultaneously occupied by Cl^-^ (Lobet and Dutzler, 2006). Coupled 2Cl^-^:H^+^ exchange arises from excursions of a highly conserved extracellular glutamate residue, E_ext_, into *S*_ext_ and *S*_cen_ anion binding sites, displacing two bound Cl ions, whereupon the carboxylate group of E_ext_ can exchange a proton with the intracellular solution (Feng et al., 2010; Feng et al., 2012). In this model the Cl^-^ transport pathway is open at both ends during transitions of E_ext_ to the extracellular solution, potentially resulting in uncoupled movement of Cl^-^; however, it is suggested that a large kinetic barrier for Cl^-^ transitions between *S*_cen_ and *S*_int_ prevents Cl^-^ diffusion in ‘open’ states (Feng et al., 2012). An alternative theory suggests that the intracellular end of the Cl^-^ transport pathway is separately gated by a strictly conserved tyrosine residue, Y_cen_, (Accardi, 2015) in response to conformational changes affecting the carboxy-terminal region of helix O (Basilio et al., 2014), however current crystal structures of CLC exchangers do not show conformational changes of Y_cen_ that support this hypothesis (Dutzler et al., 2002; Dutzler et al., 2003; Feng et al., 2010). The intracellular proton transfer pathway diverges from the Cl^-^ transport pathway at *S*_cen_ and appears to consist of a narrow protein conduit likely accommodating a single-file arrangement of intracellular waters that facilitate H^+^ transport via a ‘water-wire’ mechanism (Cheng and Coalson, 2012; Lim et al., 2012; Han et al., 2014). Both E_ext_ and Y_cen_ are conserved in CLC channels where they are repurposed as important determinants of processes that open and close (gate) the Cl^-^ conduction pathway (Dutzler et al., 2003; Bennetts and Parker, 2013).

CLC exchangers typically favor transport of NO_3_^-^>Cl^-^>Br^-^>I^-^ (Steinmeyer et al., 1995; Friedrich et al., 1999; Vanoye and George, 2002; Hebeisen et al., 2003), whereas CLC channels characteristically favor permeation of Cl^-^>Br^-^>NO_3_^-^>I^-^ (White and Miller, 1981; Thiemann et al., 1992; Steinmeyer et al., 1994; Pusch et al., 1995; Rychkov et al., 1996; Ludewig et al., 1997; Waldegger and Jentsch, 2000). Conductance (the rate of passage of anions) of ClC-0 channels reaches a minimum in mixtures of Cl^-^ and NO_3_^-^and increases with increasing preponderance of NO_3_^-^(Pusch et al., 1995). This anomalous mole fraction dependence in mixtures of Cl^-^ and NO_3_^-^is evidence of interactions between permeating species at neighboring binding sites in the channel pore (Tabcharani et al., 1993). ClC-1 channels do not show multi-ion behavior in mixtures of Cl^-^ and NO_3_^-^, however anomalous mole fraction dependence in mixtures of Cl^-^ and SCN^-^ or ClO_4_^-^shows that ClC-1 channels can also simultaneously accommodate multiple anionic species (Rychkov et al., 1998). These findings are substantiated by recent cryo-electron microscopy (cryo-EM) structures of bovine ClC-K (Park et al., 2017) and human ClC-1 Cl^-^ channels (Park and MacKinnon, 2018) that show that the canonical, exchanger-like, Cl^-^ transport pathway is preserved in CLC channels. Based on the cryo-EM structures of ClC-K and ClC-1 it has been suggested that minor structural differences in the transport pathway underpin lower kinetic barriers for Cl^-^ transitions between *S*_cen_ and *S*_int_ that allows Cl^-^ to leak through open CLC channels (Park et al., 2017; Park and MacKinnon, 2018).

Curiously, the ClC-1 structure does not show Cl^-^ occupancy of *S*_cen_ and reveals a wide tunnel, corresponding to the narrow proton transfer conduit of CLC exchangers, that bifurcates from the canonical anion transport pathway and forms a vestibule at the intracellular surface of the protein (Park and MacKinnon, 2018). Here we show that, despite being located outside of the canonical transport pathway, replacing protein residues that line the intracellular vestibule with corresponding residues of CLC exchangers fundamentally affects characteristics of the anion permeation pathway in ClC-1 channels.

## Materials and Methods

Human *ClCN1* cDNA was expressed from pCDNA 3.1(+) expression vector in HEK293T/17 cells (ATCC) by transient transfection using Fugene 6.0 reagent (Promega) according to the manufacturer’s specifications. Mutations were introduced using the Quickchange method (Stratagene) and verified by DNA sequencing in both directions. For ClC-1 V292E/S537T/H538R+M539 mutants two separately prepared constructs were examined at 24-72 hours post-transfection, however both failed to yield functional channels. Patch clamp experiments were conducted in whole-cell configuration at room temperature (296 ± 1 K) 24-48 h post transfection using an Axopatch 200B amplifier and Digidata 1322A A/D board controlled by AxoGraph X software. Currents were obtained at a 10 kHz sampling frequency and analogue filtered at 5 kHz. Currents were recorded using AxoGraph X. Offline analysis was conducted using Axograph X, Microsoft Excel 14.3.9 and GraphPad Prism 7.0.

During experiments cells were continuously superfused with standard bath solution containing (mM); NaCl (140), CsCl (4), MgCl_2_ (2), CaCl_2_ (2), TRIS (10) adjusted to pH 8.0. For anion substitution experiments equimolar NaCl was replaced with NaBr, NaNO_3_ or NaI. Rapid solution exchange was achieved using an SF-77 fast solution exchanger (Warner Instruments). The pipette (internal) solution contained (mM) Cs-glutamate (80), CsCl (40), EGTA-Na (10), HEPES (10) adjusted to pH 7.0. After achieving whole-cell configuration no less than 5 min was allowed for pipette and internal solutions to equilibrate before experiments commenced. Patch pipettes were formed with a P-97 pipette puller (Sutter) and had resistance 1.5-3.0 MΩ when filled with the above solution. Series resistance did not exceed 5 MΩ and was 85-90% compensated.

Equilibrium membrane voltages (*V*_eq_) were recorded in current-clamp mode using cells that expressed large ClC-1 mediated currents (>5nA outward currents elicited from a +100 mV voltage pulse), to avoid inaccuracies where substituting anions blocked ClC-1. Where possible, shifts in equilibrium voltage (Δ*V*_eq_) upon anion substitution were fit with a Goldmann-Hodgkin-Katz function of the form:

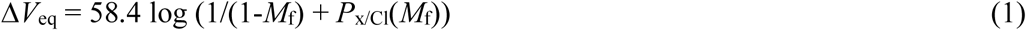

Where *M*_f_ is the mole-fraction of Cl^-^ replaced by anion X^-^ (*M*_f_ = [X^-^]/([X^-^]+[Cl^-^])) and *P*_x/Cl_ is the relative permeability of anion X^-^ with respect to Cl^-^. Channel activity was typically assessed by measuring currents in response to a +100 mV conditioning pulse for 100 ms, followed by 50 ms test pulses ranging from −120 mV to +100 mV in 20 mV increments, however in some instances test pulses ranged from between −80 mV (cells with large inward Cl^-^ currents) and −40 mV (where *V*_eq_ was shifted to more positive voltages with anion substitution) to +100 mV. Membrane voltage was clamped at −30 mv for 3 s between successive iterations. For the purpose of comparing relative conductance in mixtures of Cl^-^ and substituting anions current was measured at the end of the variable test pulse. As the ClC-1 current-voltage relationship is non-linear, outward slope conductance (corresponding to Cl^-^ and substituting anions moving in to the cell) was approximated by measuring chord conductance at a point +20 mV from the zero-current voltage (*E*_rev_), interpolated by fitting a quadratic function to the experimentally derived current-voltage relationship. For shortened voltage protocols, with less negative test pulses, no less than two data points for inward current were included in estimations of outward slope conductance. Chord conductance when I^-^ replaced Cl^-^, for S537T/H538R/T539A and S537T/H538R+M539 mutants, was calculated from the +100 mV test pulse (ie: *V* = 100 mV − *E*_rev_), as currents became outwardly rectifying in mixtures of Cl^-^ and I^-^. Relative conductance in mixtures of Cl^-^ and substituting anions was determined by normalising to conductance for the same cell in standard bath solution (152 mM Cl^-^). Liquid junction potentials were calculated using JPcalc (Barry, 1994) and corrections applied *ex post facto*.

Data points in all figures represent individual experimental determinations from *n* separate cells. Where pooled data are shown data represents mean and either SEM or 95% confidence interval, as indicated.

## Results

### The intracellular vestibule of ClC-1 channels

Human ClC-1 channels show considerable homology to the algal 2Cl^-^/H^+^ exchanger CmClC (32% identical residues and 53% sequence similarity in the membrane embedded domain). We compared the cryo-EM structure of human ClC-1 Cl^-^ channel (Park and MacKinnon, 2018) and the X-ray crystallographic structure of CmClC (Feng et al., 2010). The ClC-1 structure showed a wide intracellular vestibule diverging from the conserved Cl^-^ transport pathway in the region of the central anion binding site of ClC exchangers (Fig. 1A). In contrast the corresponding region of CmClC was confined to a narrow conduit (Fig. 1B), consistent with the proposed H^+^ transport pathway of ClC exchangers (Cheng and Coalson, 2012; Lim et al., 2012; Han et al., 2014). Both the intracellular vestibule of ClC-1 and the CmClC conduit were lined by residues that link alpha-helices O and P; S537, H538 and T539 and T474, R475 and A476, in ClC-1 and CmClC, respectively (Fig. 1 & Fig. 2). We replaced the ClC-1 helix O-P linker with the corresponding region of CmClC (ClC-1 S537T/H538R/T539A; referred to as ClC-1 CmO-P, in the interest of brevity) and examined the function of ClC-1 CmO-P chimeras using whole cell patch-clamp electrophysiology of ClC-1 expressed in HEK293T cells (Fig. 3A).

**Fig. 1.**
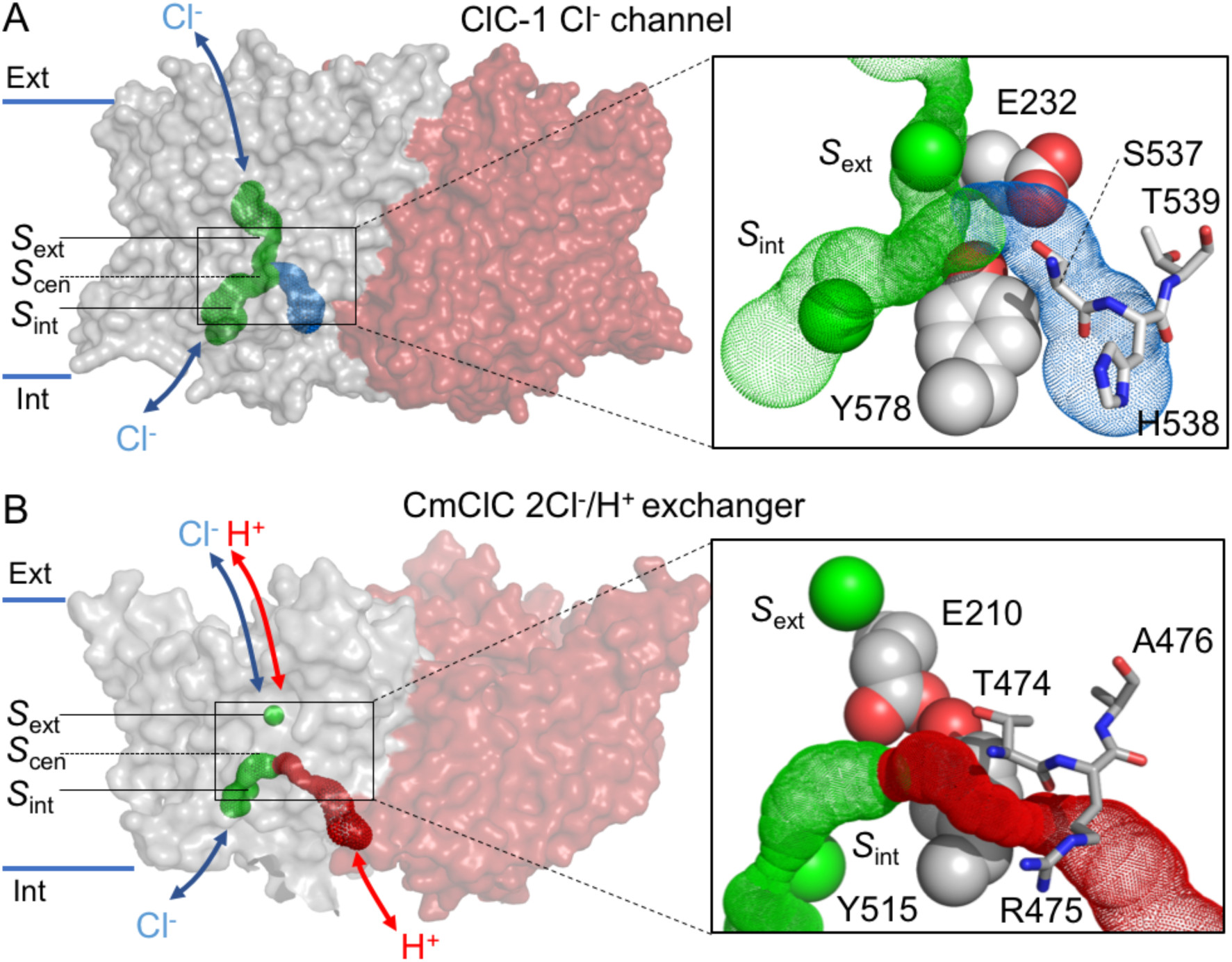
Ion transport pathways in (A) ClC −1 Cl^-^ channels and (B) CmClC 2Cl^-^/H^+^ exchangers. ClC-1 (PDB Id: 6COY) and CmClC (PDB Id: 3ORG) are depicted in as transparent surfaces colored by subunit. Approximate ion trajectories are indicated by arrows. Internal cavities corresponding to the conserved Cl^-^ transport pathway (*green*), the ClC-1 intracellular vestibule (*blue*) and the CmClC intracellular proton transport pathway (*red*) are shown as dot surfaces. Cavity detection was carried out using CAVER 3.0 (Chovancova et al., 2012). Intracellular (Int) and extracellular (Ext) protein surfaces are indicated, as are relative positions of external (*S*_ext_), central (*S*_cen_) and Internal (*S*_int_) anion binding sites. Right panels show close-up view of the boxed area in left panels. Shown are E_ext_ (ClC-1 E232, CmClC E210), Y_cen_ (ClC-1 Y578, CmClC Y515), and residues linking alpha-helices O and P (ClC-1 S537,H538,T539 and CmClC T474,R475,A476). *In situ* Cl ions are shown as *green* spheres.

**Figure 2.**
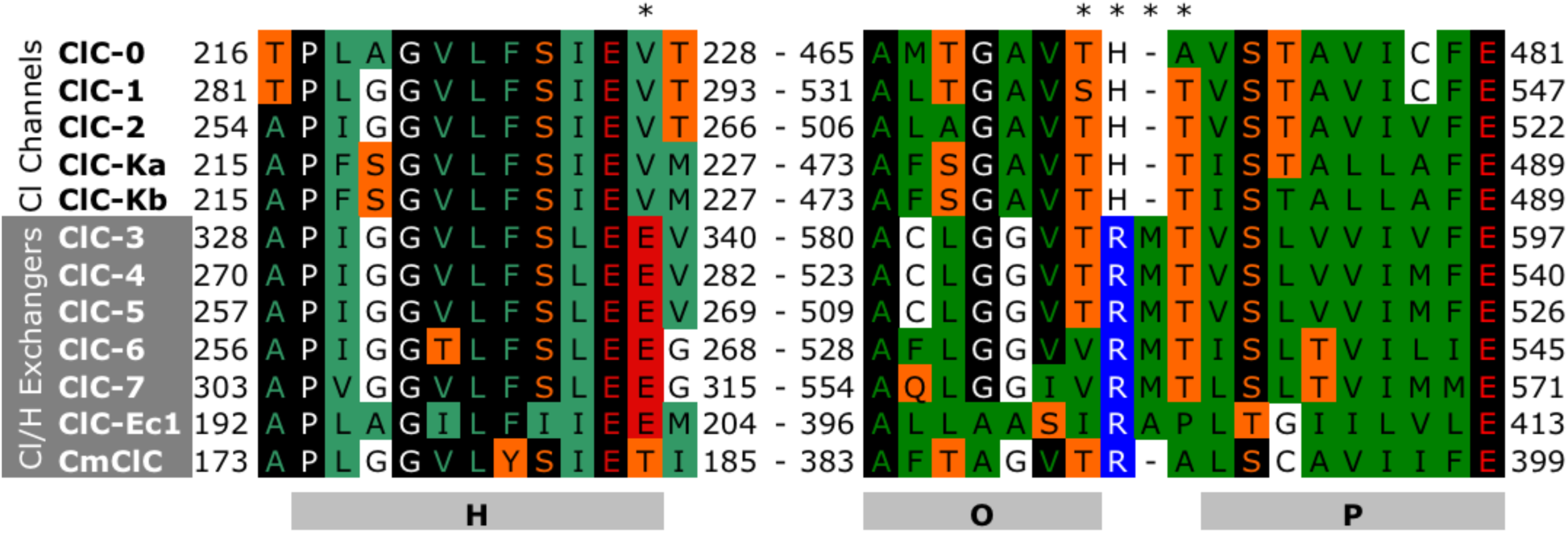
Sequence alignments for human CLC channels and exchangers in the region of helix H and the helix O-P linker. Also included are ClC-0 channel from *Torpedo marmorata*, ClC-ec1 exchanger from *Escherichia coli* and CmClC exchanger from *Cyanidioschyzon merolae*. Residues are colored according to properties; *green* non-polar, *orange* polar, *blue* basic, *red* acidic and *black* strongly conserved. * indicates ClC-1 residues mutated in the current study.

**Fig. 3.**
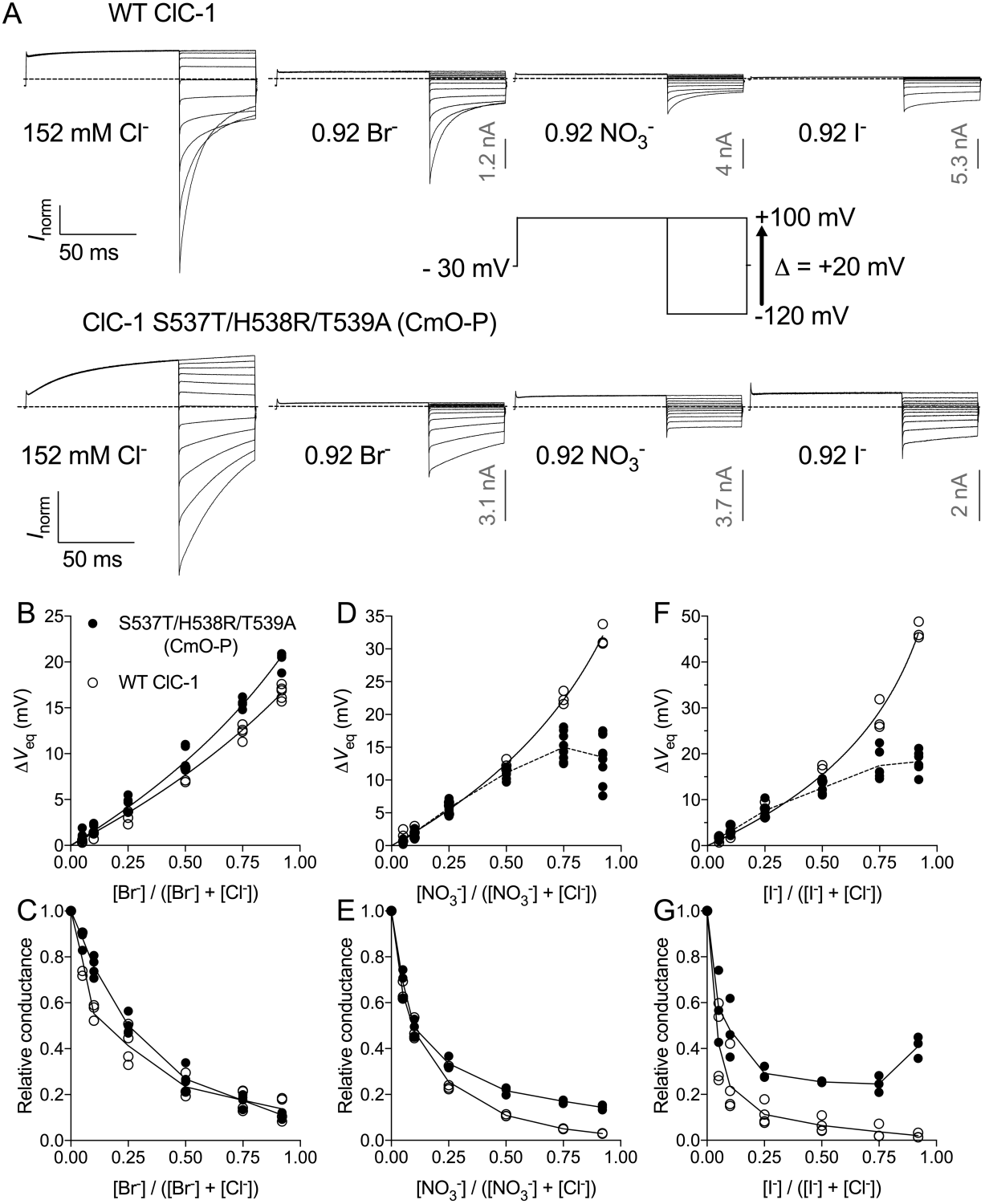
Altered anion permeation in ClC-1 S537T/H538R/T539A (CmO-P) mutants. (A) Representative whole-cell currents are shown for WT ClC-1 or the ClC-1 CmO-P mutant with 152 mM extracellular Cl^-^ (far left) or when 92% of extracellular Cl^-^ (140 mM) was substituted for Br^-^, NO_3_^-^or I^-^, as indicated. Dashed lines indicate zero-current level. Data were normalized to outward current in 152 mM Cl^-^ to compensate for cell to cell variation (that is; not all current traces were recorded from the same cell); however, scale bars (shown in gray) represent absolute current amplitudes corresponding to peak outward current determined from the same cell in 152 mM Cl^-^. Inset depicts the typical voltage protocol that was used to elicit currents. In all panels 0.3 ms of the current response following voltage steps is not shown in order to obscure transient capacitative currents. (B, D, F) Excursions of the equilibrium membrane voltage (*V*_eq_) are shown where various mole fractions of extracellular Cl^-^ were replaced with either (B) Br^-^, (D) NO_3_^-^or (F) I^-^ for WT ClC-1 (open circles) or the CmO-P chimera (filled circles). Solid lines depict fits of a Goldman-Hodgkin-Katz (GHK) equation (*Equation 1. Materials and methods*) to the experimental data points. Dotted lines connect the mean values of experimental data points where data was not adequately fit by a GHK equation. Data for anion permeability ratios and values of *n* for all mutants in this study are shown in Table 1. (C, E, G) relative conductance of ClC-1 (open circles) and CmO-P chimeras (filled circles), corresponding to permeation of ClC-1 from the extracellular to the intracellular solution, in mixtures of Cl^-^ and (C) Br^-^, (E) NO_3_^-^or (G) I^-^. Solid lines connect the mean values of experimental data points.

### Exchanger-like mutations in the ClC-1 intracellular vestibule affect anion permeation

We probed the anion permeation pathway of ClC-1 by measuring relative permeability (the ease with which an anion passes through the channel) and outward conductance (the rate of passage of anions from the extracellular to the intracellular solution) when extracellular Cl^-^ was progressively replaced by Br^-^, NO_3_^-^or I^-^. For WT ClC-1, Cl^-^ replacement resulted in predictable excursions of equilibrium membrane voltage (*V*_eq_: the membrane voltage at which there is no net ion flux) to more positive values, indicating reduced permeability of substituting anions with respect to Cl^-^ (Fig. 3B, D & F, Table 1). Concurrently, ClC-1 conductance was reduced when increasing mole fractions of Cl^-^ were replaced by less permeant anions (Fig. 3C, E & G). Our experiments were in agreement with previous studies of ClC-1 anion selectivity (Fahlke et al., 1997a; Rychkov et al., 1998), and showed that ClC-1 selects for Cl^-^>Br^-^>NO_3_^-^>I^-^ (Table 1), and that Cl^-^ and Br^-^, NO_3_^-^or I^-^ permeate ClC-1 without appreciably interacting with one another inside the channel pore. In contrast, CmO-P chimeras showed anomalous mole-fraction dependence in mixtures of Cl^-^ and NO_3_^-^or I^-^ (Fig. 3D, F & G). Increasing mole fractions of Cl^-^ replacement by NO_3_^-^or I^-^ resulted in increasing permeability ratios for substituting anions (Fig. 4). The anion selectivity sequence of CmO-P channels was Cl^-^>NO_3_^-^∼I^-^∼Br^-^ (Table 1). Similarly, CmO-P conductance was increased, with respect to WT ClC-1, in mixtures of Cl^-^ and NO_3_^-^or I^-^ (Fig. 3E & G); and conductance reached a minimum when 50-75% of Cl^-^ was replaced by I^-^ (Fig. 3G). Anomalous mole-fraction dependence of CmO-P permeability in mixtures of Cl^-^ and NO_3_^-^or I^-^ showed interactions between permeating anionic species at consecutive binding sites in CmO-P mutants (Tabcharani et al., 1993).

**Table 1.** Permeability ratios (*P*_X_/*P*_Cl_) for WT ClC-1 and mutants were determined from fits of *equation 1* to experimental data points. Where mixtures of anions resulted in **\***anomalous mole-fraction dependence *P*_X_/*P*_Cl_ is shown for 92% replacement of Cl^-^. Data are means and SEM determined from *n* separate experiments.

| | $P_{Br}/P_{Cl}$ | $n$ | $P_{NO_3}/P_{Cl}$ | $n$ | $P_I/P_{Cl}$ | $n$ | Permeability sequence |
| --- | --- | --- | --- | --- | --- | --- | --- |
| WT ClC-1 | $0.48 \pm 0.01$ | 5 | $0.22 \pm 0.01$ | 3 | $0.09 \pm 0.01$ | 3 | $Cl^- > Br^- > NO_3^- > I^-$ |
| ClC-1 CmO-P | $0.40 \pm 0.01$ | 4 | **\*$0.54 \pm 0.10$** | 10 | **\*$0.44 \pm 0.05$** | 6 | $Cl^- > NO_3^- \sim I^- \sim Br^-$ |
| S537T/H538R+M539 | - | | **\*$0.56 \pm 0.04$** | 3 | **\*$0.48 \pm 0.03$** | 3 | $Cl^- > NO_3^- \sim I^-$ |
| ClC-1 V292E | - | | $0.27 \pm 0.01$ | 5 | **\*$0.27 \pm 0.02$** | 4 | $Cl^- > I^- \sim NO_3^-$ |
| CmO-P/Y578A | - | | $0.26 \pm 0.01$ | 4 | $0.18 \pm 0.01$ | 4 | $Cl^- > NO_3^- > I^-$ |
| CmO-P/E232A | - | | **\*$0.62 \pm 0.06$** | 4 | $0.24 \pm 0.01$ | 3 | $Cl^- > NO_3^- > I^-$ |

**Fig. 4.**
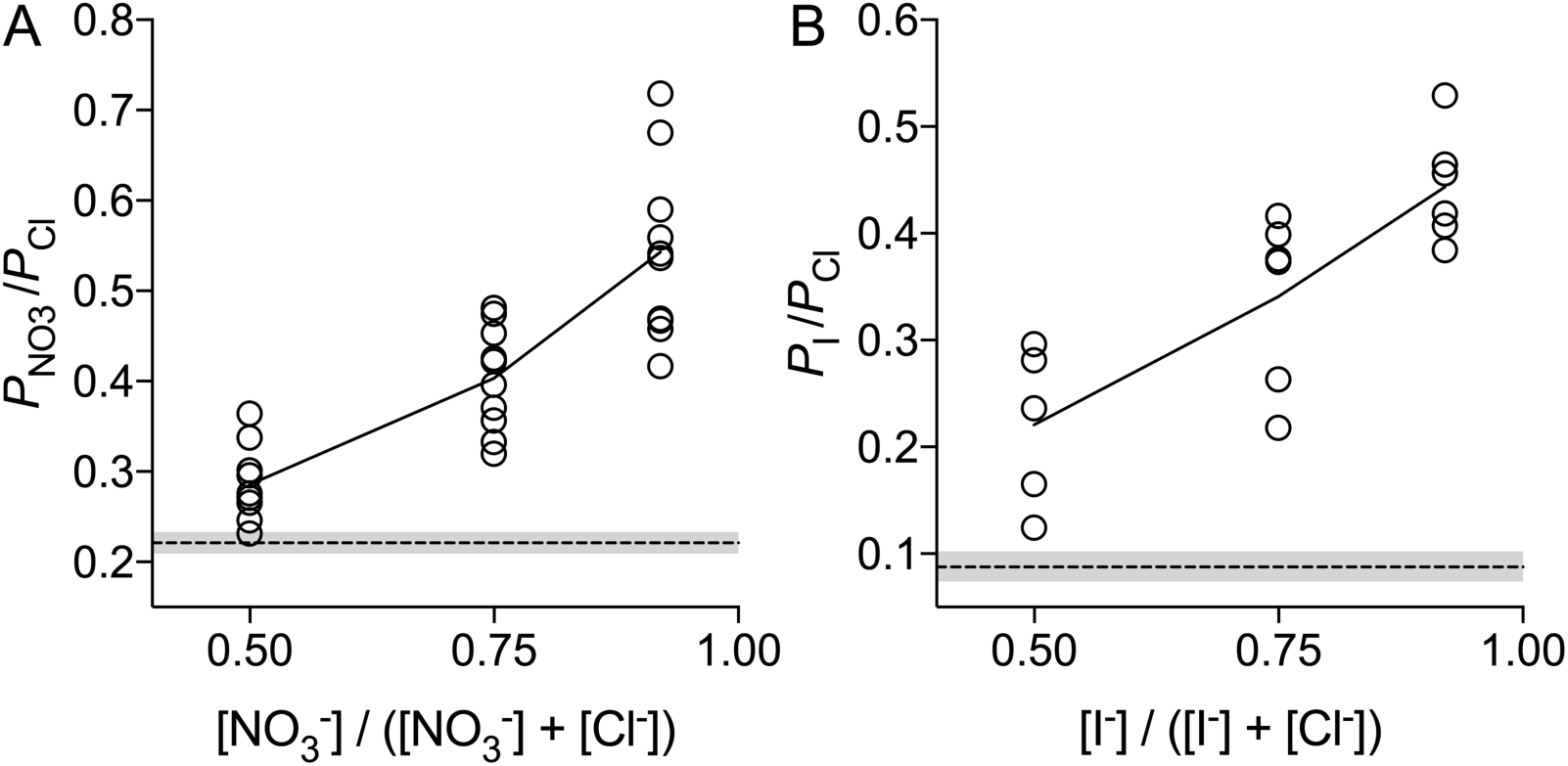
Anomalous mole-fraction dependence of anion permeability in CmO-P chimeras. Permeability ratios (*P*_(X)_/*P*_Cl_) for ClC-1 CmO-P mutants were determined from fits of *equation 1* to individual data points shown in Fig.3D & F at discrete mole-fractions of Cl^-^ replacement by (A) NO_3_^-^(*n* = 10) or (B) I^-^ (*n* = 6). Solid lines connect the mean of permeability ratios determined for the CmO-P chimera. Dashed lines and shaded area represent permeability ratios and 95% confidence intervals, respectively, determined from fits of *equation 1* to experimental data points for WT ClC-1; shown in Fig.3D & F.

Human CLC exchangers are more distantly related to CLC channels than algal CmClC exchangers and, currently, no three-dimensional structures of human CLC exchangers have been resolved. All human CLC exchangers have an extra methionine residue in the region linking helices O and P that is not present in CLC channels or in CmClC exchangers (Fig. 2). We replaced the region linking helices O and P of ClC-1 with a typical human-like exchanger sequence (S537T/H538R+M539). Our experiments showed that ClC-1 S537T/H538R+M539 chimeras displayed comparable multi-ion pore behaviour in mixtures of Cl^-^ and NO_3_^-^or I^-^ to CmO-P mutants (Fig. 5 and Table 1). Human exchangers also have a conserved glutamate residue corresponding to the intracellular vestibule residue V292 of ClC-1 that, while not conserved in CmClC, appears to be a critical determinant of the proton transport pathway for most CLC exchangers (Accardi et al., 2005; Zdebik et al., 2008; Lim and Miller, 2009). ClC-1/V292E mutants showed similar behaviour to WT ClC-1 in mixtures of Cl^-^ and NO_3_^-^(Fig. 6C & D), however; equilibrium potentials for V292E mutants showed anomalous mole fraction dependence in mixtures of Cl^-^ and I^-^ (Fig. 6E). In contrast to WT ClC-1 (Cl^-^>NO_3_^-^>I^-^) V292E mutants displayed a Cl^-^>I^-^=NO_3_^-^selectivity sequence (Table 1).

**Fig. 5.**
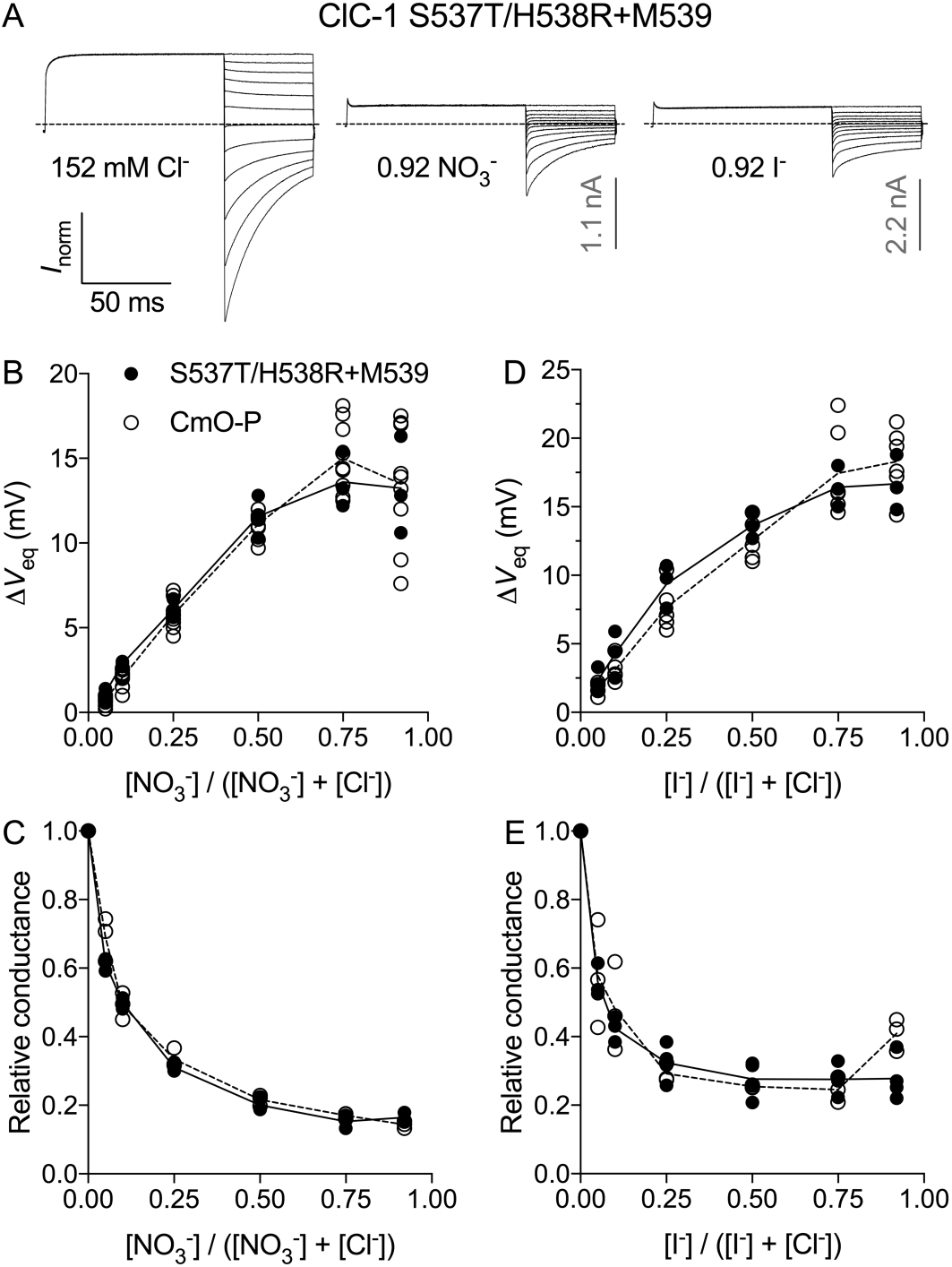
Altered anion permeation in human-exchanger like ClC-1 S537T/H538R + M539 mutants. (A) Representative whole-cell currents are shown for WT ClC-1 or ClC-1 mutant with 152 mM extracellular Cl^-^ (far left) or when 92% of extracellular Cl^-^ (140 mM) was substituted for NO_3_^-^or I^-^, as indicated. Dashed lines indicate zero-current level. Data were normalized to peak outward current in 152 mM Cl^-^ to compensate for cell to cell variation. Scale bars (*grey*) represent peak outward current in 152 mM Cl^-^. The voltage protocol used to elicit currents was as shown in Fig. 3A. In all panels 0.3 ms of the current response following voltage steps is not shown in order to obscure transient capacitative currents. (B, D) Excursions of the equilibrium membrane voltage (*V*_eq_) are shown where various mole fractions of extracellular Cl^-^ were replaced with either (B) NO_3_^-^or (D) I^-^ for CmO-P (S537T/H538R/T539A) (open circles) or S537T/H538R+M539 mutant (filled circles). Solid lines depict fits of a Goldman-Hodgkin-Katz (GHK) equation (*Equation 1. Materials and methods*) to the experimental data points. Dotted lines connect the mean values of experimental data points where data was not adequately fit by a GHK equation. Data for anion permeability ratios and values of *n* are shown in Table 1. (C, E) relative conductance of CmO-P (open circles) and S537T/H538R+M539 mutants (filled circles), corresponding to permeation from the extracellular to the intracellular solution, in mixtures of Cl^-^ and (C) NO_3_^-^or (E) I^-^. Solid lines connect the mean values of experimental data points for S537T/H538R+M539 mutants, dotted lines connect means for CmO-P mutants.

**Fig. 6.**
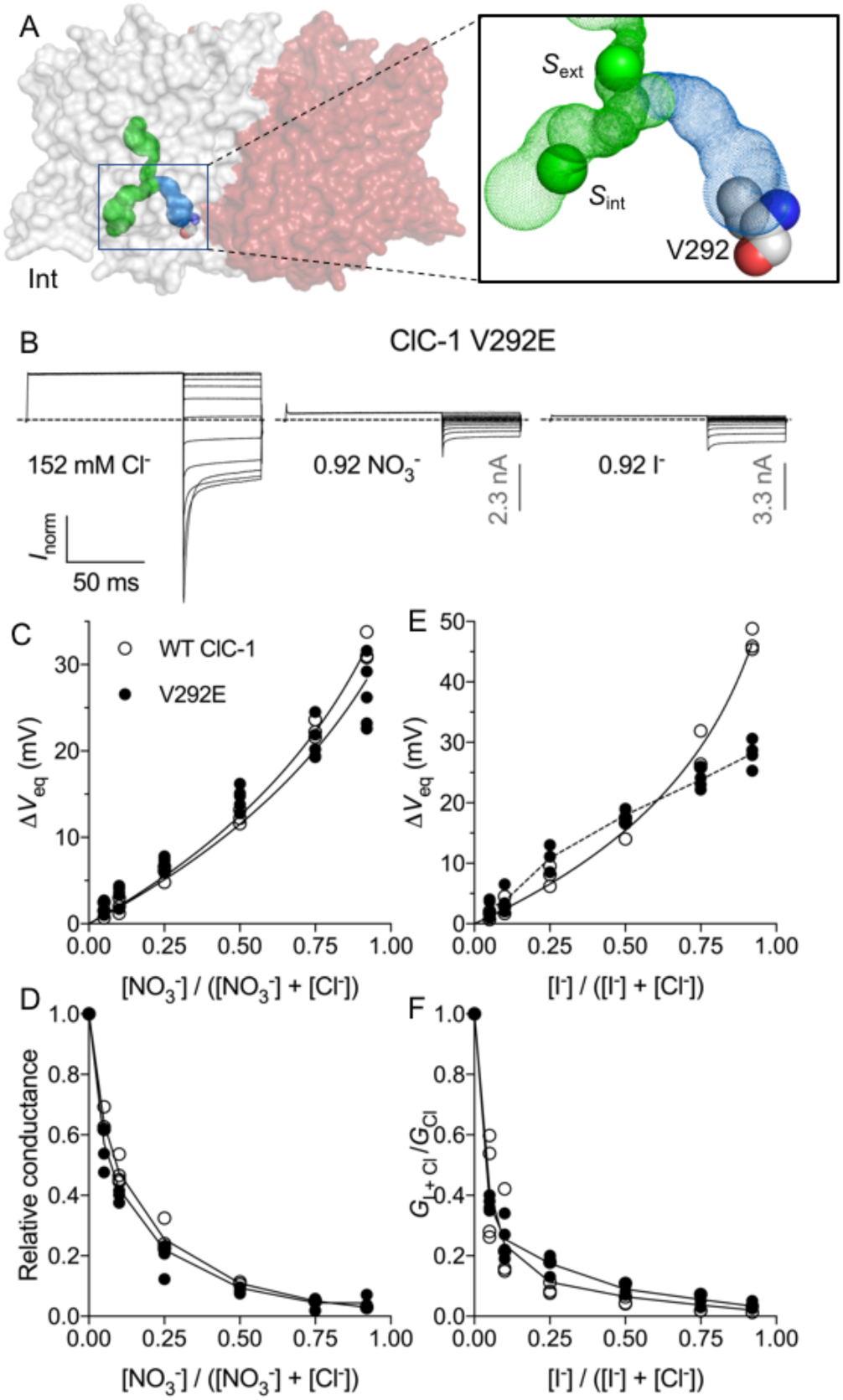
Altered I^-^ permeability in ClC-1/V292E mutants. (A) The ClC-1 cryo-EM structure (PDB id: 6COY) is depicted in transparent surface format, colored by subunit. Inset shows close-up view of the boxed area, with internal cavities depicted as dot surfaces, and residue V292 depicted in space-fill format. *In situ* Cl ions bound to the external (*S*_ext_) and internal (*S*_int_) sites are shown as green spheres. (B) Representative current traces obtained from ClC-1/V292E mutants in 152 mM extracellular Cl^-^ or when 92% of extracellular Cl^-^ (140 mM) was substituted NO_3_^-^or I^-^, as indicated. Dashed lines indicate zero-current level. The voltage protocol used to elicit currents was as shown in Fig. 3A. Data were normalized to peak outward current in 152 mM Cl^-^. Scale bars (*grey*) represent peak outward current in 152 mM Cl^-^. In all panels 0.3 ms of the current response following voltage steps is not shown in order to obscure transient capacitative currents. (C, E) Excursions of the equilibrium membrane voltage (*V*_eq_) are shown for experiments where various mole-fractions of extracellular Cl^-^ were replaced by either (C) NO_3_^-^or (E) I^-^ for ClC-1 (open circles) or CmO-P/V292E mutants (filled circles). Dashed lines connect mean values of data points for V292E mutants in mixtures of Cl and I ions, solid lines depict fits of *equation 1* to the experimental data points. Relative permeability ratios determined from these fits and values of *n* are shown in Table 1. (D, F) Relative conductance of ClC-1 (open circles) and ClC-1/V292E mutants (filled circles) in mixtures of Cl^-^ and NO_3_^-^(D) or I^-^ (F). Solid lines connect mean values of experimental data points.

Although ClC-1 S537T/H538R+M539 chimeras (Fig. 5) and ClC-1/V292E mutants (Fig. 6) robustly expressed functional channels, combined ClC-1 V292E/S537T/H538R+M529 mutants were not functional (not shown); suggesting either defective protein folding, or that mutants were locked in a non-conductive conformation.

### Removing the side-chain of Y_cen_ removes multi-ion behaviour in ClC-1 CmO-P mutants

In CLC exchanger structures (Dutzler et al., 2002; Dutzler et al., 2003; Feng et al., 2010), and the ClC-K channel structure (Park et al., 2017) the phenolic-hydroxyl group of Y_cen_ is shown to coordinate Cl^-^ at the central anion binding site, or alternatively, the carboxylic acid group of E_ext_ in the CmClC structure (Feng et al., 2010). In the ClC-1 channel structure no occupancy of *S*_cen_ is observed and Y_cen_ (Y578 in ClC-1) is displaced, with respect to CLC exchanger structures, by ∼1.5 Å; away from the conserved Cl^-^ transport pathway toward the intracellular vestibule (Park and MacKinnon, 2018). Y_cen_ mutations have little effect on anion permeation in ClC channels (Accardi and Pusch, 2003; Chen and Chen, 2003; Estevez et al., 2003), however; Y_cen_ mutations allow leakage of Cl^-^ in ClC-exchangers (Accardi et al., 2006), where the number of Cl^-^ that leak during an H^+^ exchange half-cycle is inversely proportional to the volume of the substituent side chain (Walden et al., 2007). Additionally, mutation of Y_cen_ in ClC-ec1 exchangers removes Cl^-^ occupancy of *S*_cen_ in crystal structures (Accardi et al., 2006) by greatly reducing Cl^-^ binding affinity (Picollo et al., 2009). Our results showed that exchanger-like mutations in the ClC-1 intracellular vestibule resulted in anomalous mole-fraction dependence in mixtures of Cl^-^ and NO_3_^-^or I^-^, that is indicative of interactions between anions bound to neighbouring sites in the permeation pathway (Figs 3-5). One possibility is that intracellular vestibule mutations could affect the distribution of anions in the conserved transport pathway by altering the conformation of Y_cen_ in ClC-1 CmO-P chimeras. Consistent with this hypothesis CmO-P/Y578A mutants did not show anomalous mole-fraction dependence in mixtures of Cl^-^ and NO_3_^-^or I^-^ (Fig. 7) and displayed similar Cl^-^>NO_3_^-^>I^-^ selectivity to WT ClC-1, although with increased relative I^-^ permeability (*P*_I/Cl_ = 0.09 ± 0.01 and 0.18 ± 0.01 for WT ClC-1 and CmO-P/Y578A, respectively) (Table 1).

**Fig. 7.**
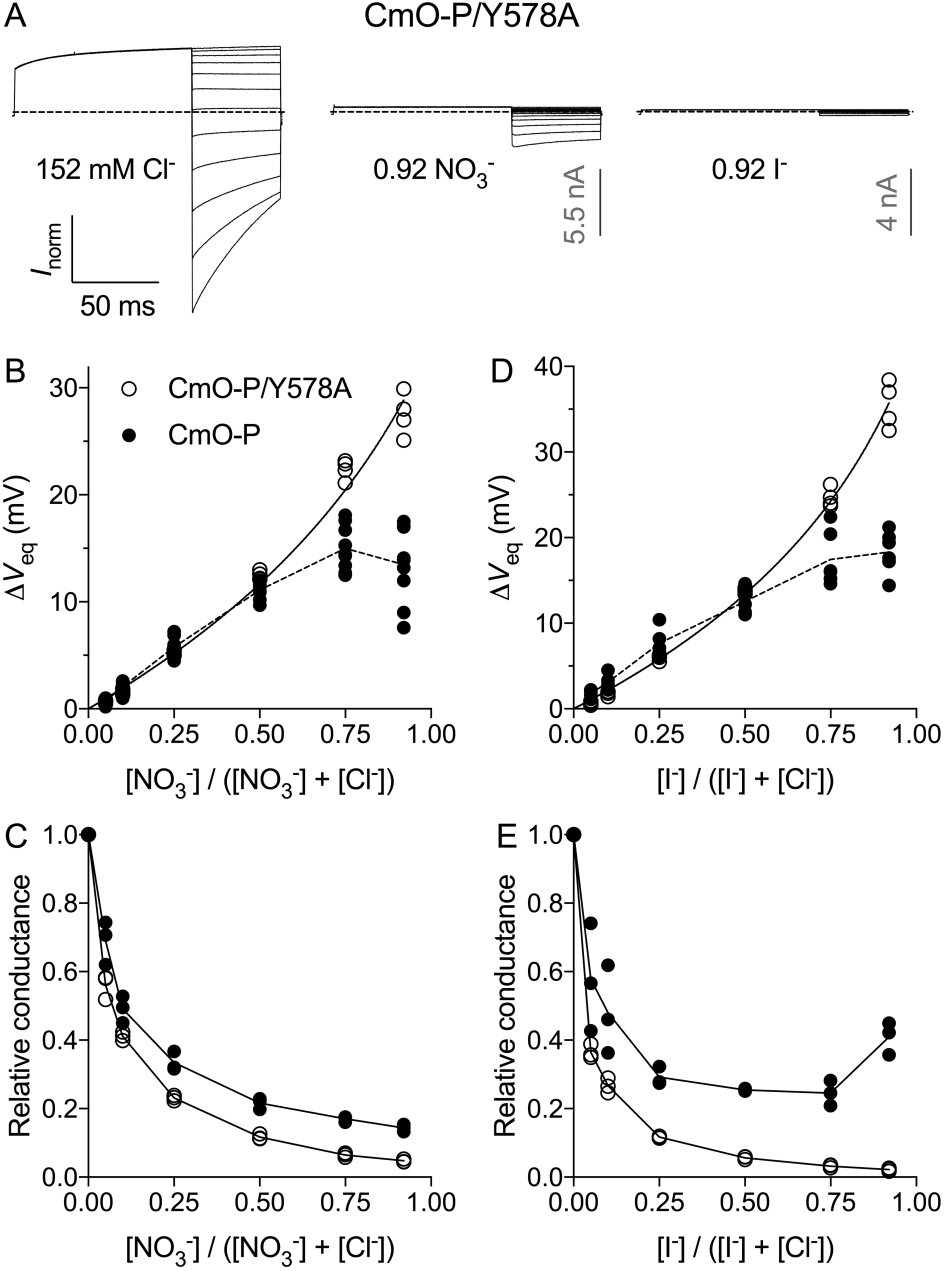
Removal of the aromatic side-chain of Y578 removes multi-ion behaviour of CmO-P mutants. (A) Representative current traces obtained from CmO-P/Y578A mutants in 152 mM extracellular Cl^-^ or when 92% of extracellular Cl^-^ (140 mM) was substituted NO_3_^-^or I^-^, as indicated. Dashed lines indicate zero-current level. The voltage protocol used to elicit currents was as shown in Fig. 3A; except for 92% I^-^ replacement, where test voltages ranged from −40 to +100 mV, as detailed in *Materials and Methods*. Data were normalized to peak outward current in 152 mM Cl^-^. Scale bars (*grey*) represent peak outward current in 152 mM Cl^-^. In all panels 0.3 ms of the current response following voltage steps is not shown in order to obscure transient capacitative currents. (B, D) Excursions of the equilibrium membrane voltage (*V*_eq_) are shown for experiments where various mole-fractions of extracellular Cl^-^ were replaced by either (B) NO_3_^-^or (D) I^-^ for ClC-1 CmO-P (filled circles) or CmO-P/Y578A mutants (open circles). Dashed lines connect mean values of data points for CmO-P, solid lines depict fits of a Goldman-Hodgkin-Katz (GHK) equation (*Equation 1. Materials and methods*) to the experimental data points for CmO-P/Y578A. Relative permeability ratios determined from these fits and values of *n* are shown in Table 1. (C, E) Relative conductance of CmO-P (filled circles) and CmO-P/Y578A mutants (open circles) in mixtures of Cl^-^ and NO_3_^-^(D) or I^-^ (F). Solid lines connect mean values of experimental data points.

### Removing the side-chain of the gating glutamate, E_ext_, fails to remove multi-ion behaviour in ClC-1 CmO-P chimeras

Similar to its role in shuttling protons through the outer and middle Cl^-^ binding sites of CLC exchangers, block of CLC channels by the carboxylic acid side-chain of E_ext_ is responsible for gating of CLC channels. The structure of ClC-1 shows a novel conformation of E_ext_ (E232 in ClC-1) where the side-chain is oriented away from the Cl^-^ transport pathway towards the intracellular vestibule (Fig. 1A) suggesting that the structure represents an open state of ClC-1 (Park and MacKinnon, 2018). Mutations of E_ext_ have minor effects on anion permeability of ClC-1 (Fahlke et al., 1997b), however it is possible that mutations in the ClC-1 intracellular vestibule could affect the conformation of E_ext_ and affect the distribution of anions in the conserved transport pathway. While WT ClC-1 (Fig. 2A), ClC-1/E232Q (Fahlke et al., 1997b), and ClC-1 intracellular vestibule mutants (Figs 3A, 5A & 6A) all displayed inward current rectification, ClC-1 CmO-P/E232A mutants showed Cl^-^ mediated currents that weakly rectified in both outward and inward directions at high voltages (Fig. 8A). Both permeability (Fig. 8B) and outward conductance (Fig. 8C) of CmO-P/E232A mutants displayed anomalous mole-fraction dependence in mixtures of Cl^-^ and NO_3_^-^and at high mole-fractions of NO_3_^-^(92% replacement of Cl^-^ by NO_3_^-^) conductance was greater than in Cl^-^ (Fig 8C). In contrast to ClC-1 CmO-P chimeras, CmO-P/E232A mutants did not show anomalous mole-fraction dependence in mixtures of Cl^-^ and I^-^ (Fig. 8D & E). In terms of anion permeability, the selectivity sequence for CmO-P/E232A mutants was similar to WT ClC-1 (Cl^-^>NO_3_^-^>I^-^), although with increased relative permeability of NO_3_^-^and I^-^ (Table 1.). Alternatively, conductance of CmO-P/E232A mutants appeared to favour NO_3_^-^>Cl^-^>I^-^ (Fig. 8).

**Figure 8.**
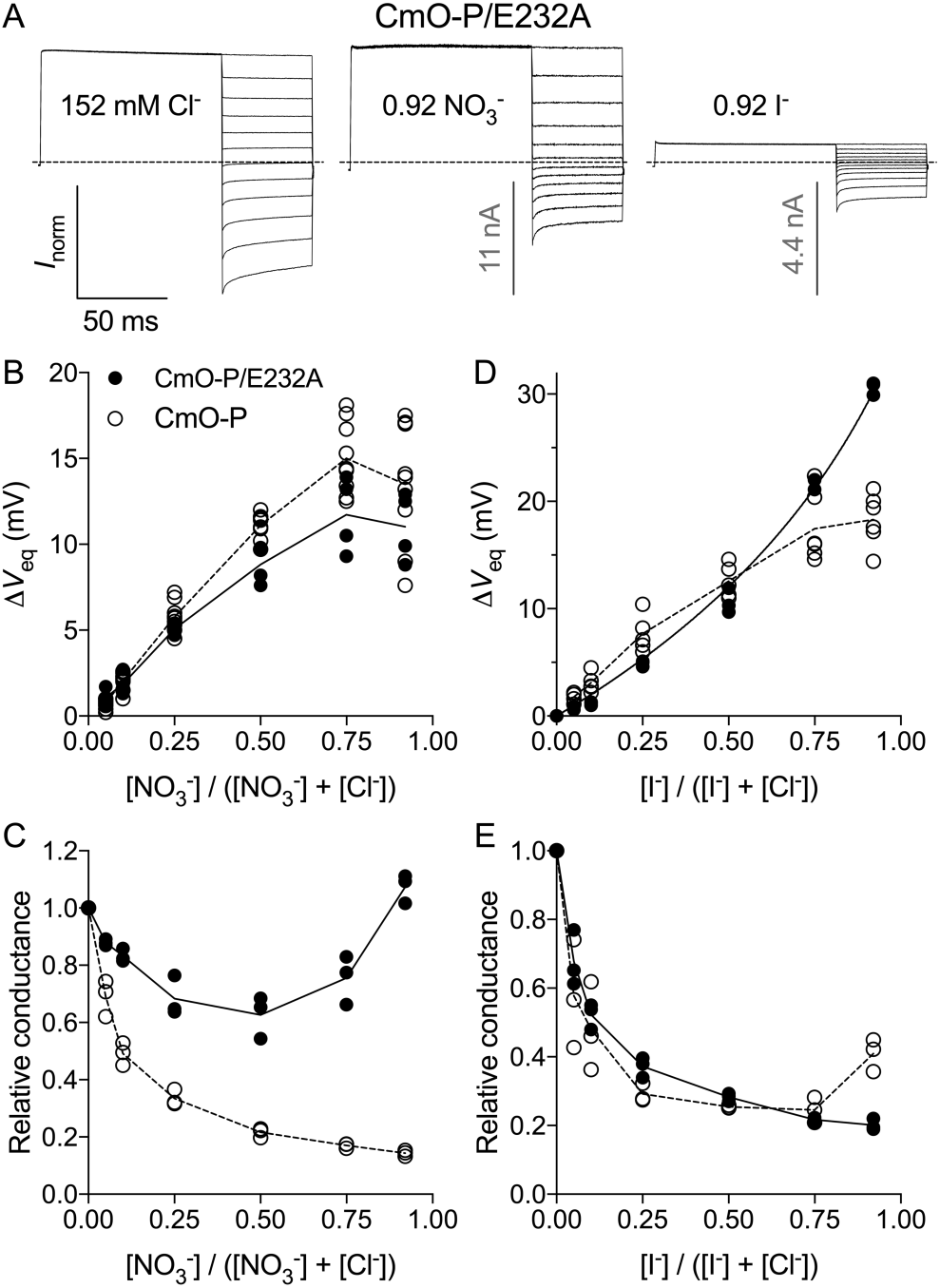
Multi-ion behavior of CmO-P/E232A mutants in mixtures of Cl^-^ and NO_3_^-^. (A) Representative current traces obtained from CmO-P/E232A mutants in 152 mM extracellular Cl^-^ or when 92% of extracellular Cl^-^ (140 mM) was substituted NO_3_^-^or I^-^, as indicated. Dashed lines indicate zero-current level. The voltage protocol used to elicit currents was as shown in Fig. 3A. Data were normalized to peak outward current in 152 mM Cl^-^. Scale bars (*grey*) represent peak outward current in 152 mM Cl^-^. In all panels 0.3 ms of the current response following voltage steps is not shown in order to obscure transient capacitative currents. (B, D) Excursions of the equilibrium membrane voltage (*V*_eq_) are shown for experiments where various mole-fractions of extracellular Cl^-^ were replaced by either (B) NO_3_^-^or (D) I^-^ for ClC-1 CmO-P or CmO-P/E232A mutants. Dashed lines connect mean values of data points for CmO-P, solid lines depict fits of a Goldman-Hodgkin-Katz (GHK) equation (*Equation 1. Materials and methods*) to the experimental data points for CmO-P/Y578A. Relative permeability ratios determined from these fits and values of *n* are shown in Table 1. (C, E) Relative conductance of CmO-P (filled circles) and CmO-P/Y578A mutants (open circles) in mixtures of Cl^-^ and NO_3_^-^(D) or I^-^ (F). Solid lines connect mean values of experimental data points.

## Discussion

For more than a decade researchers have puzzled at how, despite closely related architecture, some CLC proteins function as dissipative Cl^-^ channels and others as ion-exchangers that couple transport of Cl^-^ to counter-transport of protons (Miller, 2006). Recent high-resolution structures of CLC channels have highlighted this puzzle by showing that both functional classes of CLC proteins share a strongly conserved Cl^-^ transport pathway (Park et al., 2017; Park and MacKinnon, 2018). Current theories suggest that the difference between CLC channels and exchangers lies in subtle structural differences in the conserved pathway that allow Cl^-^ to ‘leak’ through CLC channels (Park et al., 2017; Park and MacKinnon, 2018). Our findings show that relatively minor structural differences between CLC channels and exchangers, in a recently identified intracellular vestibule, fundamentally affect the anion permeation pathway of ClC-1 channels.

The recent cryo-EM structures of bovine ClC-K (Park et al., 2017) and human CLC-1 channels (Park and MacKinnon, 2018) show that the intracellular part of the conserved Cl^-^ permeation pathway is wider in CLC channels than in CLC exchangers, supporting the theory that kinetic barriers limiting the rate of Cl^-^ transitions between binding sites are smaller in CLC channels. The structural differences that lead to lowered kinetic barriers in the region of *S*_cen_ – *S*_int_ are different in ClC-K and ClC-1, however. In ClC-K channels the orientation of a strongly conserved serine residue in the protein loop linking alpha-helices C-D (S189 in ClC-1), near *S*_cen_, is altered with respect to other CLC structures, leading to a wider pathway between *S*_cen_ – *S*_int_ (Park et al., 2017). In contrast, smaller side-chains of residues lining this region (T475 and G483 in ClC-1) are proposed to result in lower kinetic barriers in ClC-1 (Park and MacKinnon, 2018). In addition, the ClC-1 structure does not show occupancy of *S*_cen_ suggesting that this site has a low affinity for Cl^-^ (Park and MacKinnon, 2018). The ClC-1 structure also shows that displacement of Y_cen_, away from *S*_cen_, by ∼ 1.5 Å compared to other CLC structures, contributes to the wider dimensions of the Cl^-^ transport pathway in this region and to the generous dimensions of the inlet to the intracellular vestibule that bifurcates from the conserved pathway. The ClC-1 intracellular vestibule is wide, and polar, enough to accommodate Cl^-^ (diameter ∼ 3.6 Å) and intracellular water molecules (diameter ∼ 2.8 Å). Our findings show that replacing residues that line the ClC-1 intracellular vestibule with residues from CLC exchangers altered anion permeation of ClC-1; in the main part, by introducing interactions between permeating anions at neighbouring sites in the permeation pathway. These findings indicate that the mutations alter the distribution of anions in the conserved pathway, suggesting two possibilities.

First, it is likely that exchanger-like mutations in the ClC-1 intracellular vestibule allosterically affect the conformation of residues in the conserved Cl^-^ permeation pathway. A number of scenarios could account for this hypothesis however the unique conformation of residues E_ext_ and Y_cen_ in the ClC-1 structure (Park and MacKinnon, 2018), in addition to their proximity to the intracellular vestibule (Fig. 1A), suggest the possible involvement of these residues. Several studies have suggested that movements of Y_cen_ affect the Cl^-^ transport pathway of CLC exchangers (Jayaram et al., 2008; Basilio et al., 2014; Khantwal et al., 2016). During the transport cycle of prokaryotic ClCec1 exchangers it appears that conformational changes in helix O are communicated to Y_cen_ by the residue corresponding to S537 of ClC-1 (ClC-ec1 I402), and that Y_cen_ potentially acts as an intracellular gate of the transport pathway (Basilio et al., 2014). Conformational changes in helix P also occur during the transport cycle of ClC-ec1 exchangers (Khantwal et al., 2016). One possibility is that mutations in the helix O-P linker region of ClC-1 affect the distribution of anions in the conserved pathway by altering the conformation of Y_cen._ This hypothesis may account for our results by an altered conformation of Y_cen_ promoting anion occupancy of ClC-1 *S*_cen_, similar to that seen in structures of CLC exchangers (Dutzler et al., 2002; Dutzler et al., 2003) and ClC-K channels (Park et al., 2017), in turn leading to interactions between permeating anions at neighbouring binding sites. This hypothesis is supported by removal of multi-ion effects by removing the aromatic side-chain of Y_cen_ in CmO-P/Y578A mutants (Fig. 7). However, while the crystal structure of ClC-ec1 shows that I402 (equivalent to ClC-1 S537) directly contacts Y_cen_ (Dutzler et al., 2002) the equivalent residues in ClC-1 are separated by ∼ 5 Å (Park and MacKinnon, 2018). Movements of E_ext_ in the canonical transport pathway are intrinsic to the proton transport mechanism of CLC exchangers (Dutzler et al., 2002; Dutzler et al., 2003; Feng et al., 2010; Feng et al., 2012) and gating of CLC channels (Dutzler et al., 2003), however the ClC-1 structure shows a novel conformation of E_ext,_ oriented away from the canonical pathway towards the intracellular vestibule (Fig. 1A). It is also possible that exchanger-like mutations in the ClC-1 intracellular vestibule could affect the canonical pathway by altering the conformation of E_ext_; although the closest approach that the mutations make to E_ext_ is ∼ 8 Å. To some extent this hypothesis is supported by the removal of multi-ion behaviour in mixtures of Cl^-^ and I^-^ (Fig. 8D), however CmO-P/E232A mutants showed strong multi-ion behaviour in mixtures of Cl^-^ and NO_3_^-^(Fig. 8B & C). Alternatively, ClC-1 V292E mutants displayed multi-ion effects in mixtures of Cl^-^ and I^-^ (Fig. 6D), although it is not readily apparent how the V292E mutation could affect the affect the conformation of Y_cen_ or E_ext_, as these side-chains are separated by ∼ 10 Å and 9 Å, respectively, and lie on opposite sides of the intracellular vestibule (Park and MacKinnon, 2018).

The second possibility is that, as suggested by Park and MacKinnon (Park and MacKinnon, 2018), the ClC-1 intracellular vestibule may function as a secondary pore that allows permeating anions to escape from the canonical pathway. Replacing residues that line the inlet of the ClC-1 intracellular vestibule with the corresponding residues of CLC exchangers possibly pinches off the inlet (Fig. 1), leading to anion permeation via consecutive binding sites in the canonical pathway. The narrowest region of the ClC-1 intracellular vestibule and the CmClC conduit is at the junction with the conserved Cl^-^ transport pathway, and in both instances the side-chain of Y_cen_ forms part of this junction (Fig. 1). Our experiments showed that removing the side-chain of Y_cen_ removed multi-ion behaviour and restored WT ClC-1 anion selectivity to CmO-P chimeras (Fig. 5), possibly by allowing permeating anions to leak in to the intracellular vestibule. Mutation V292E, near the intracellular mouth of the ClC-1 vestibule also altered ClC-1 anion selectivity by introducing multi-ion effects in mixtures of Cl^-^ and I^-^ (Fig. 6). It is possible that electrostatic repulsion between the carboxylic acid group of E292 and relatively large I^-^ (diameter 4.4 Å) could favour permeation via the conserved pathway, leading to anomalous mole-fraction effects in mixtures of Cl^-^ and I^-^. Similarly, removal of a negative charge at the inlet of the vestibule in CmO-P/E232A mutants may affect I^-^ permeation in CmO-P chimeras. It is worth noting that ClC-1 channels are blocked by intracellular application of 9-anthracene carboxylic acid (9-AC) and, intriguingly, mutation of residues S537 and H538 practically removes 9-AC block, and most residues identified by Estevez and co-authors that strongly affect ClC-1 inhibition by 9-AC cluster around the intracellular vestibule (Estevez et al., 2003). In addition, removal of the side-chain of E_ext_ increases the blocking affinity of anionic 9-AC in ClC-1 (Estevez et al., 2003) and another anionic intracellular inhibitor, *p*-chlorophenoxyacetic acid (CPA), in ClC-1 (Estevez et al., 2003) and ClC-0 channels (Traverso et al., 2003). These findings suggest the possibility that block of CLC channels by 9-AC and CPA could arise from binding in the intracellular vestibule.

The ClC-1 intracellular vestibule bifurcates from the conserved Cl^-^ transport pathway near the middle of the protein and appears to have evolved from distension of a narrow protein conduit that supports proton transport in CLC exchangers. It is likely that exchanger-like mutations in this region have allosteric effects on the Cl^-^ transport pathway of ClC-1 as poorly understood conformational changes in this region occur during gating of CLC channels (Accardi and Pusch, 2003) and during the transport cycle of CLC exchangers (Basilio et al., 2014; Khantwal et al., 2016). Our results also suggest the possibility that the intracellular vestibule of ClC-1 could function as a secondary pore that allows permeating anions to escape from the canonical transport pathway. The ClC-1 structure appears to be representative of an open state (Park and MacKinnon, 2018); ultimately, structures of closed and intermediate states of ClC-1 will likely be required to elucidate how structural changes in the intracellular vestibule affect the anion permeation pathway. While current theories suggest that subtle differences in the canonical Cl^-^ transport pathway account for the disparate molecular mechanism of CLC channels and exchangers our findings show that structural differences in the recently identified intracellular vestibule of ClC-1 channels strongly affect the Cl^-^ permeation pathway.

## Author contribution

Brett Bennetts conceived the study, constructed mutants, conducted electrophysiological experiments and data analysis and wrote the original manuscript. Craig Morton and Michael Parker analysed and advised on structural data. All authors edited the final manuscript.

## Footnotes

a The name of the CLC superfamily arises from an acronym of **Cl** ion **C**hannel; a vestige from the original, long held, belief that all CLC proteins were chloride ion channels. In this paper we have adopted the contemporary nomenclature (Jentsch and Pusch, 2018), where the CLC family is generically referred to in upper-case (for example, CLC channels) and individual proteins are referred to using their historic designations, with a lower-case ‘l’ (for example, ClC-1).

